# Complex phyllosphere microbiome aids in the establishment of the invasive macrophyte *Hydrilla verticillata* (L.) under conditions of nitrogen scarcity

**DOI:** 10.1101/2021.01.11.426196

**Authors:** Jiří Bárta, Caio Cesar Pires de Paula, Eliška Rejmánková, Qiang Lin, Iva Kohoutová, Dagmara Sirová

## Abstract

Despite the low availability of nitrogen (N), the highly productive macrophyte *Hydrilla verticillata* (L.) is a successful invader of the littoral zones at lake Atitlán, Guatemala, with profound implications for lake ecology. To help answer the question of how *Hydrilla*, accompanied by the filamentous green alga *Cladophora* Kützing (Ulvophyceae), sustains fast growth under conditions of N scarcity, we studied the composition and potential biogeochemical function of the associated microbiomes. We combined results from next generation sequencing of associated bacterial and fungal assemblages with traditional microscopy-based taxonomical evaluation of algae and cyanobacteria. We focused on the presence of specific N_2_-fixing genera and their relative importance. Data on community composition are complemented with measurements of diazotrophic activity. The results expand our knowledge of the ecophysiology of these algae-plant-microbe consortia and suggest that several levels of biological complexity should be considered to fully understand aquatic plant ecology and the process of macrophyte invasions.

## Introduction

Aquatic macrophytes are important in structuring aquatic environments, promoting increased habitat complexity at different scales and supporting the functional diversity of other assemblages, including invertebrates and fish. Therefore, invasions of lake littoral zones by adventive macrophyte species can facilitate major cascading changes in their ecology [1]. Central American lakes, specifically volcanic lakes located in the Central American Volcanic Arc, have been largely unexplored by invasion biologists and the information on the impact of macrophyte invasions on this type of ecosystem is limited [2].

Lake Atitlán in the highlands of western Guatemala is one of the most important waterbodies in Central America. It is a nitrogen (N) limited hard-water lake, whose trophic state has recently transitioned from oligotrophic to mesotrophic [3]. The littoral zone covers about 4% of the lake area and is now dominated by the introduced *Hydrilla veriticillata* (Hydrocharitaceae), a plant native to the warm regions of Asia. This submersed macrophyte is one of the most successful aquatic invaders with a wide ecological amplitude, fast growth rates, and high dispersion ability. *Hydrilla* therefore has a great potential to invade a variety of habitats, often resulting in important physical, chemical, and biotic effects on the freshwater environments worldwide [4,5]. These changes are not always considered solely negative, however, and the ecological services *Hydrilla* provides need to be considered [6] *Hydrilla* possesses some remarkable ecophysiological traits: it can grow at depths exceeding 10 m, near the water surface it branches profusely to form a dense canopy intercepting light to the exclusion of other submersed plants [7], it is photosynthetically highly efficient, may utilize bicarbonate as a carbon source if the lake pH and carbonate concentrations are high, and, during the night, it can also switch to C4-like carbon metabolism [8]. The dense foliage breaking the water surface often supports vigorous growth of green filamentous algae such as *Cladophora* spp. [2]. The presence of *Hydrilla* in the lake was first detected in 2002 and it has since substantially changed the littoral biogeochemistry, such as water column dissolved oxygen profiles and macroinvertebrate diversity [2]. Despite the low N availability, the highly productive *Hydrilla* populations at Atitlán were shown to have lower C:N:P ratios than native species, contributing to faster decomposition and overall fast biomass turnover rates [2]. The questions which intrigued us were: How does *Hydrilla* sustain such fast growth under low N availability? Can phyllosphere-associated microbes in *Hydrilla* contribute to the success of the species by supplementing some of the host demands for N? Do they differ in composition and function to those found on the *Hydrilla* companion, *Cladophora*?

An important ecophysiological aspect in some macrophytes, observed more frequently in submerged species with small root:shoot ratios and simple anatomy, including *Hydrilla*, is the reduced role of the rhizosphere [9,10]. In these species, the leaf and stem surfaces and tissues become the main interfaces for carbon exudation [11] and nutrient absorption, and provide a diversity of microbial niches that can harbor microorganisms with a broad spectrum of metabolic capabilities (Ariosa et al. 2004, Pettit et al. 2016, Mathai et al. 2019). It is now becoming widely recognized that plant-associated microorganisms, both endophytic and epiphytic, can become important functional drivers of their eukaryotic hosts [12]. Aquatic macrophytes, as well as macroscopic filamentous algae, often host unique assemblages of microbiota on their surfaces and within their tissues, which differ significantly from those found in the plankton [11,13]. These complex microbial communities likely influence many functions of their host and contribute to the nutrient cycling and energy flow in aquatic ecosystem [14,15]. Examples of potential microbial activities include N_2_ fixation, nitrification and denitrification, Fe (III) reduction; Fe (II) oxidation, sulfate reduction, methanogenesis, methanotrophy, and many others. Nitrogen fixation has previously been found associated with several macrophyte species [16–18] as well as seagrasses [19,20]. Epiphytic microbiota have also been investigated on large filamentous algae such as *Cladophora* spp. and shown to display potential for biogeochemical cycling and traits that foster host growth [21–23]. It follows that the metabolic roles of macrophytes in lakes, the associated organic matter production, nutrient cycling, as well as their invasiveness should not be viewed separate from their microbiomes. Studies that address biological invasions of macrophytes in freshwater ecosystems from the point of view of “holobionts” - assemblages of hosts and the many other species living in or around them, which together form discrete ecological units [24], have so far been very rare [25].

In order to fill some of these knowledge gaps and tackle the question of whether the microbial communities associated with both *Hydrilla* and *Cladophora* can contribute to N cycling in the littoral zones of N-limited lake Atitlán, we evaluated the following: (1) Differences in microbial assemblages associated with *Hydrilla* and *Cladophora* in terms of composition and potential function; (2) the effect of sampling location on *Hydrilla* and *Cladophora* microbiomes (3) the N_2_-fixation activity of the *Hydrilla* and *Cladophora* holobionts and its correlation with the presence of N_2_-fixers. We used a combination of next generation amplicon sequencing and nitrogenase activity measurements to gain new insights on the potential role of algae-plant-microbe ecological interactions in lake littoral zones.

## Material and Methods

### Study site and field sampling

Lake Atitlán is one of numerous volcanic lakes located in the Central American Volcanic Arc in the highlands of western Guatemala; for its detailed description see Rejmánková et al. 2018b. The littoral zone (water depth of ≤ 10 m with vascular plants present) covers only about 4.8 km^2^, which is < 4% of the total lake area [26]. Historically, the littoral was dominated by a bulrush, *Schoenoplectus californicus*, and deeper areas by a diverse group of submersed macrophytes (e.g. *Potamogeton illinoensis*, *P. pectinatus*, *Ceratophyllum demersum*, *Chara* spp.). During the last decade, the composition of the macrophyte flora has changed quite dramatically due to the introduction of *H. verticillata*. A massive development of green algae (mostly from the genus *Cladophora*) and a dense periphyton of dominant diatoms (genera *Cymbella*, *Gomphoneis*, *Epithemia*, *Nitzschia*) on stems and leaves of littoral vegetation occur commonly at sites near numerous inflows of sewage water. The switch from low-productive littoral zones dominated by emergent *S. californicus* to high-productive littoral zones dominated by submersed *Hydrilla* represents a major acceleration of biogeochemical cycling in these lake zones [2].

We sampled littoral areas in bays near the San Lucas, Santiago, and San Juan settlements in April/ May 2016 (Fig. S1); each site included an area of roughly 1 ha ranging from 1m to 8 m depth and dominated by *Hydrilla*. *Hydrilla* and *Cladophora* leaf and stem samples were collected in three random replicates from the canopy at each site and placed inside sterile Whirl-Pak bags. Water column samples were collected into sterile Nalgene bottles. Samples of *Cladophora* mats were gently collected from the *Hydrilla* canopy and samples of *Hydrilla* were collected from stands which did not contain any visible *Cladophora* mats. At San Juan, there was only a limited area of *Hydrilla* with attached *Cladophora* filaments and thus we don’t have *Cladophora* replicates from this site. We also conducted limited sampling of *Hydrilla* and *Cladophora* holobionts in July 2016 and of the native macrophyte species *Potamogeton illinoensis* and *P. pectinatus* holobionts in May 2018 for reference measurements of N_2_-fixation activity. *Azolla filiculoides* Lam. (Azollaceae) samples were included as positive control in the N_2_-fixation activity assay. This aquatic fern forms a symbiotic relationship with the highly efficient N_2_-fixing cyanobacterium *Anabaena azollae* (Nostocaceae). Macrophyte biomass was assessed as described in [27]. Subsamples of *Hydrilla* and *Cladophora* holobionts were preserved in formaldehyde (2% v/v) and later used for microscopic evaluation of cyanobacteria.

All samples were transported to the laboratory in a cooler within 2-3 h of collection. Water quality parameters including temperature, dissolved oxygen concentration, and pH were recorded with hand-held YSI 556 meter at each site (Table 1). Water samples collected from the upper 0—20 cm layer were analyzed for inorganic N (NH_4_-N, NO_3_-N), soluble reactive phosphorus (SRP), total N, and total P. Water samples for NO_3_–N, NH_4_–N, and SRP were filtered through a 0.45 µm filter within an hour following sampling and frozen until analysis. Nitrogen species and total N were analyzed on a Lachat 8000 (Hach Company, Loveland, CO, USA) flow injection analyzer using method # 10-107-04-1-B (cadmium column reduction), method # 10-107-06-1-F (indophenol), and a modified method # 10-115-01-4-F (persulfate digestion) for NO_3_–N, NH_4_–N, and total N, respectively. Soluble reactive P was analyzed by the ascorbic acid method of Murphy & Riley (1962).

**Table 1.** Means (n = 3) and standard deviations (SD) of water nutrients (μg L^−1^), the biomass of *Hydrilla* canopy (g of dry mass m^−2^), and percent cover in littoral zones of Lake Atitlan, April 2016, for locations see Fig. 1 in Supplement. Lower biomass and cover at Santiago site is due to a recent *Hydrilla* removal by local residents to allow boat passage.

| 785 Location | 786 Depth Range<br>(m) | NH<sub>4</sub>-N |  | NO<sub>3</sub>-N |  | SRP |  | TN |  | TP |  | Biomass |  | % cover |
| --- | --- | --- | --- | --- | --- | --- | --- | --- | --- | --- | --- | --- | --- | --- |
|  |  | Mean | SD | Mean | SD | Mean | SD | Mean | SD | Mean | SD | Mean | SD |  |
| 787 |  |  |  |  |  |  |  |  |  |  |  |  |  |  |
| 788 San Lucas | 1 - 8 | 7.0 | 5.5 | 3.7 | 4.2 | 15.2 | 7.7 | 537 | 115 | 81 | 27 | 1838 | 1168 | 90 |
| 789 Santiago | 1 - 6 | 10.3 | 6.3 | 7.6 | 5.5 | 6.8 | 2.1 | 242 | 78 | 38 | 7 | 908 | 755 | 60 |
| 790 San Juan | 1 - 4 | 5.7 | 5.2 | 5.1 | 7.6 | 14.7 | 9.5 | 389 | 111 | 57 | 20 | 1152 | 504 | 90 |
| 791 |  |  |  |  |  |  |  |  |  |  |  |  |  |  |
| 792 Lake center | ~ 300 | 2.7 | 3.2 | 2.2 | 1.9 | 6.1 | 3.7 | 188 | 60 | 32 | 11 | - |  | - |
| 793 |  |  |  |  |  |  |  |  |  |  |  |  |  |  |

### Nitrogenase activity assays

Leafy branches from *Hydrilla* canopy, and the *P. illinoensis* and *P. pectinatus* native plants, were collected from three random spots at each location and transported to the field laboratory in small buckets with lake water. After gently shaking off the excess water, about 10 g of fresh plant material, equivalent to 1.5-2 g dry weight (DW) of *Hydrilla* or *Cladophora* with their respective epiphyton, was transferred into sterile 275 mL glass jars with 200 mL of filtered lake water. A preliminary test was conducted to verify that this sample manipulation did not impact nitrogenase activity. In the test we performed the acetylene reduction assay (ARA) [29] in samples treated as described above with those that were collected and immediately placed to the fixation vials where the oxygen was lowered by exchange of part of the headspace for N_2_ gas. There were no significant differences in the nitrogenase activity measured by ARA between the two treatments over 2, 5, 10, 18, and 25 hours of incubation (t-test, P = 0.7; data not shown). The acetylene reduction technique, ARA (Stal 1988), was employed to estimate N_2_-fixation activity via the reduction of acetylene to ethylene by nitrogenase. Ten percent of the headspace were replaced with acetylene gas, freshly generated from calcium carbide, and the jars were incubated for 4 hours for daytime measurements (10am – 2pm) and 8 hours for nighttime measurements (9pm – 5am). At the end of the exposure, 7-8 mL of headspace was withdrawn with an airtight syringe (Alltech) and analyzed by gas chromatograph (Shimadzu 14 GC) with a flame ionization detector and a Porapak-T column at 800°C. The results are reported as the nitrogenase activity in nmol C_2_H_4_ g of plant dry weight (DW)^−1^ h^−1^. Controls run as samples without acetylene addition as well as blanks (tubes without plant tissue incubated with acetylene) showed no signs of endogenous ethylene production. After terminating the exposure, samples were divided, half was kept for dry weight determination and half was freeze-dried for subsequent DNA extraction. To determine potential carbon limitation in the heterotrophic N_2_-fixers during the night, we added glucose to night exposures of subsamples of both *Hydrilla* and *Cladophora* holobionts. The differences between “glucose plus” and “glucose minus” treatments were not significant (data not shown).

### DNA extraction and amplicon sequencing

Approximately 0.5 g FW of *Hydrilla* phyllosphere/*Cladophora* filament sample was added to a FastPrep^TM^ Lysis Matrix E tube (MP Biomedicals, Solon, OH, USA). Hexadecyltrimethylammonium bromide (CTAB) extraction buffer, containing 5% CTAB (in 0.7 M NaCl, 120 mM potassium phosphate, pH 8.0) and 0.5 ml phenol-chloroform-isoamylalcohol (25:24:1), was added and agitated in a FastPrep Instrument (MP Biomedicals, Solon, OH, USA) at speed 5–6 for 45 s. After bead beating, the samples were extracted with chloroform and precipitated in a PEG 6000/1.6 M NaCl solution. Pellets were washed with 70% ethanol and re-suspended in molecular biology grade water. Total DNA was quantified using known concentration of genomic DNA of *E.coli* which was used for creation of calibration curve and, after addition of fluorescent dye SybrGreen, the fluorescent signal was compared with unknown samples [30].

### Sequence data analysis

The aliquots of DNA extracts were sent to SEQme company (Prague, Czech republic) for the preparation of a library and sequencing using MiSeq platform. The Earth Microbiome Project (EMP) protocol was used for library preparation with modified universal primers 515FB/806RB45 and ITS1F/ITS246 for prokaryotic 16S rDNA and fungal ITS1 amplicons, respectively. The coverage of prokaryotic primer pair 515FB/806RB was additionally tested in-silico using ARB Silva database release 132. The primer pair 515FB/806RB covers almost uniformly all major bacterial and archaeal phyla [31]. Both bacterial 16SrDNA raw pair-end reads (150 bp) were joined using ea-utils to obtain reads of approx. 250 bp length [32]. Quality filtering of reads was applied as previously described [33]. After quality filtering the sequences were trimmed to uniform length 250 bp. Before picking the operational taxonomic units (OTU), the fungal ITS1 region was extracted from reads using ITSx algorithm [34]. USEARCH 8.1 was used for OTU table construction with the following parameters: similarity cut-off 97% and 98.5% for prokaryotes and fungi, respectively, singletons were discarded. Taxonomy was assigned with BLAST algorithm and SILVA 132 [35] and UNITE 7.2 [36] databases were used for annotation in case of bacteria and fungi, respectively. Raw sequences of 16SrDNA and ITS1 amplicons were deposited in the European Nucleotide Archive (ENA) under the study ID PRJEB42360.

### Microbial co-occurence analysis

To better understand the interactions among the bacterial and fungal communities, we constructed the ecological networks by calculating all possible Spearman correlation coefficients between the different OTUs. Spearman’s Rho between the pairwise OTUs matrix was constructed using the “Hmisc” R package. The false discovery rate (FDR) controlling procedure was used to calculate the P-values for multiple testing [37]. A valid co-occurrence was considered robust if the Spearman correlation coefficient was either equal to or greater than 0.6 or −0.6 and statistically significant if P-values <0.01. The cut-off correlation of 0.6 or −0.6 was chosen to increase the confidence in the validity of the interactions. Network images were generated in R with the help of the “Igraph” package. We used the undirected network (where the edge has no direction) and the Fruchterman–Reingold layout.

### Statistical treatment

To characterize a set of microbes consistently present in the hosts (core microbiome), we used a detection threshold of 0.01% and a prevalence threshold of 99.9% (i.e., a given taxon must be present in 99.9% of samples of the respective host with a relative abundance of at least 0.01%) using the microbiomeR package [38]. Richness, Shannon-Weiner Index, and Evenness were calculated using the *BiodiversityR* package [39]. To test whether the environmental conditions (host or location) differ significantly in the microbiome structure, we performed a one-way permutational multivariate analysis of variance (PERMANOVA). PERMANOVAs were based on Euclidean distance matrix with 999 permutations using the *adonis* function in the *vegan* package. Pearson's correlation was used to verify the relationship between the abundance of N_2_-fixers and nitrogenase activities, considering variables positively correlated with r ≥0.90. Significant differences in microbiome taxonomic composition between the two hosts were determined by linear discriminant analysis (LDA) effect size (LEfSe: http://huttenhower.sph.harvard.edu/galaxy/) [40]. The threshold on the logarithmic LDA score for discriminative features was set at 2.0 and α = 0.05, and features with at least 2.0 log-fold changes were considered significant. LEfSe detects differentially distributed lineages with the Kruskall-Wallis test, and then checks the consistency of subclass distinctions with the pairwise Wilcoxon text. The final linear discriminant analysis was used to rank all differentiating lineages by their effect size. The functional predictions were identified using the functional gene pipeline & repository (FunGene) database [41] and the Student’s t tests with a 5% probability threshold was used to verify the significance of the differences among the hosts. All graphs were generated using Paleontological Statistics software (PAST) [42] and R version 3.6.1 (R Core Team: www.R-project.org).

## Results

### Water characteristics and plant biomass

The nutrient content in the water column of the lake littoral zone is approximately double compared to that in the pelagial for both the inorganic and total forms of N and P (Table 1). The three littoral locations did not differ in their nutrient content except for the total N which was significantly higher in San Lucas than in Santiago (Table 1). The average biomass of *Hydrilla* did not differ significantly across the sampling sites due to relatively large spatial variability, and averaged 1838, 908, and 1152 g DW m^−2^.

### Microbiome composition

The final dataset consisted of 1,169,538 and 618,444 total sequences for bacteria and fungi, respectively. The average per sample was 12,712 16S rDNA sequences and 6,722 ITS sequences. After taxonomy assignment, a total of 2,899 bacterial and 910 fungal OTUs (Operational Taxonomic Units) were detected. Bacterial assemblages consisted of 29 phyla; *Hydrilla* and *Cladophora* hosted similar bacterial taxa at the class level (Fig. 1): *Bacilli*, *Gammaproteobacteria*, and *Alphaproteobacteria* dominated the assemblages; the anaerobic *Clostridia*, and *Oxyphotobacteria* (Cyanophyceae), *Planctomycetacia*, and *Bacteroidia* were present at smaller relative proportions of the total OTUs. Fungi dataset showed ten different phyla (*Ascomycota*, *Basidiomycota*, *Blastocladiomycota*, *Chytridiomycota*, *Entomophthoromycota*, *Glomeromycota*, *Mortierellomycota*, *Mucoromycota*, *Olpidiomycota*, and *Rozellomycota*) and 25 classes, with an additional 5.8% of OTUs unclassified at the class level. Fungal assemblages associated with the two hosts were also similar at the class level (Fig. 1), with *Dothideomycetes*, *Microbotryomycetes*, *Eurotiomycetes*, and *Leotiomycetes* as the main groups identified.

**Figure 1.**
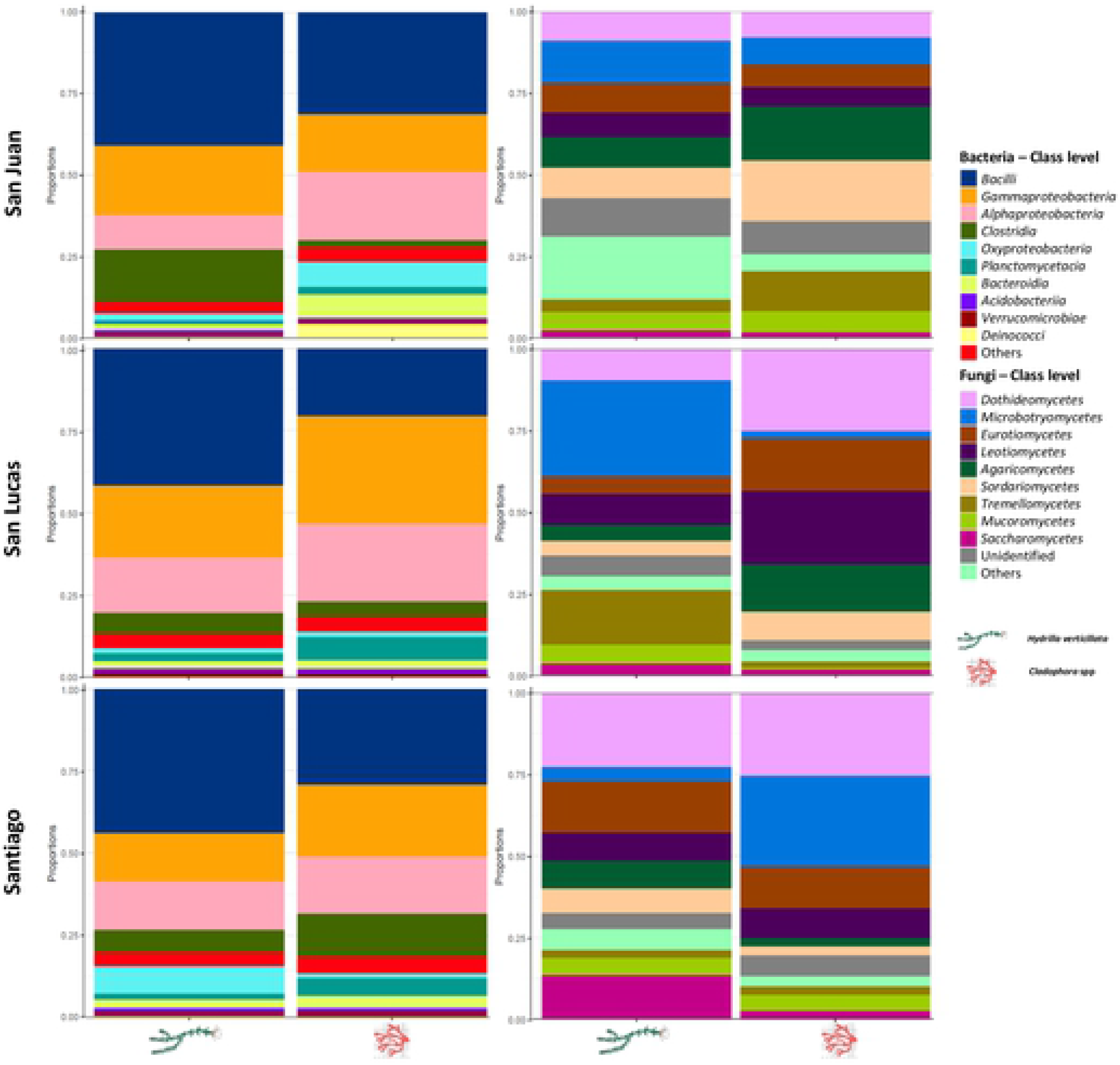
Taxonomic profiles with the most abundant taxa (bacteria and fungi) in *Hydrilla* and *Cladophora* microbiomes from San Juan, San Lucas, and Santiago locations.

Microbiome (bacteria and fungi) species richness (Chao1) did not differ significantly (p > 0.1) among the three sampled locations, nor between *Cladophora* and *Hydrilla* hosts (Fig. 2). On the other hand, microbial diversity (Fig. 2) was higher in *Cladophora* samples compared to *Hydrilla*, however these assemblages were also significantly more heterogeneous (Simpson, Fig. 2). Location (PERMANOVA, F = 6.00, p = 0.1) did not play a significant role in shaping the diversity and eveness patterns in the studied assemblages. However, the host identity (PERMANOVA, F = 10.36, p = 0.001) was a significant driver in microbiome structuring.

**Figure 2.**
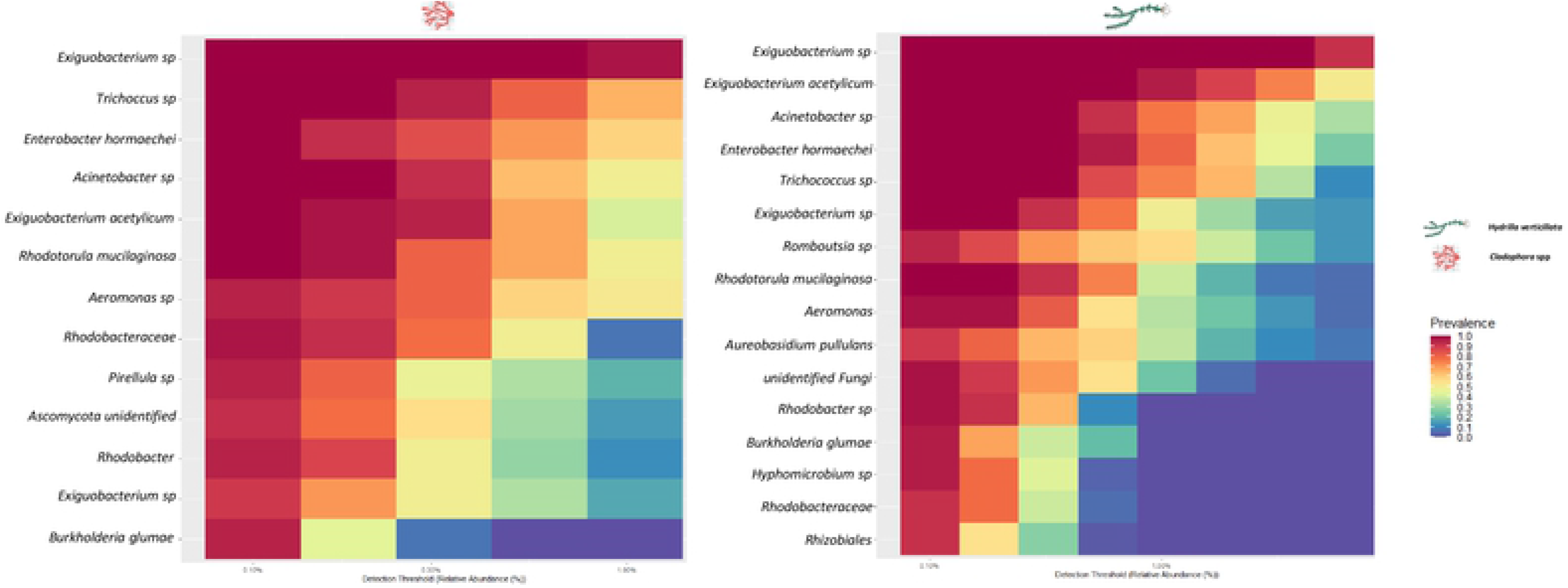
Prevalence chart of core microorganisms (bacteria and fungi) in *Cladophora* and *Hydrilla* microbiomes. All taxa shown are present in 99.9% of samples with a relative abundance of at least 0.01%.

The microbiomes of the two hosts also shared many core taxa at the genus level (Fig.3): the most important core bacterial genera in both *Cladophora* and *Hydrilla* hosts were *Exiguobacterium* (Bacilli), *Trichococcus* (Bacilli), *Pirellula* (Planktomycetes), *Hyphomicrobium* (Alphaproteobacteria), *Phreatobacter* (Alphaproteobacteria), and *Enterobacter* (Gammaproteobacteria); the unicellular yeast *Rhodotorula* (Basidiomycota) was a shared core fungal taxon. There were also core microbial taxa found to be associated with only one host: for example the *Hydrilla*-only core microbiome, included the fungal genus *Aureobasidium* (Ascomycota).

**Figure 1.**
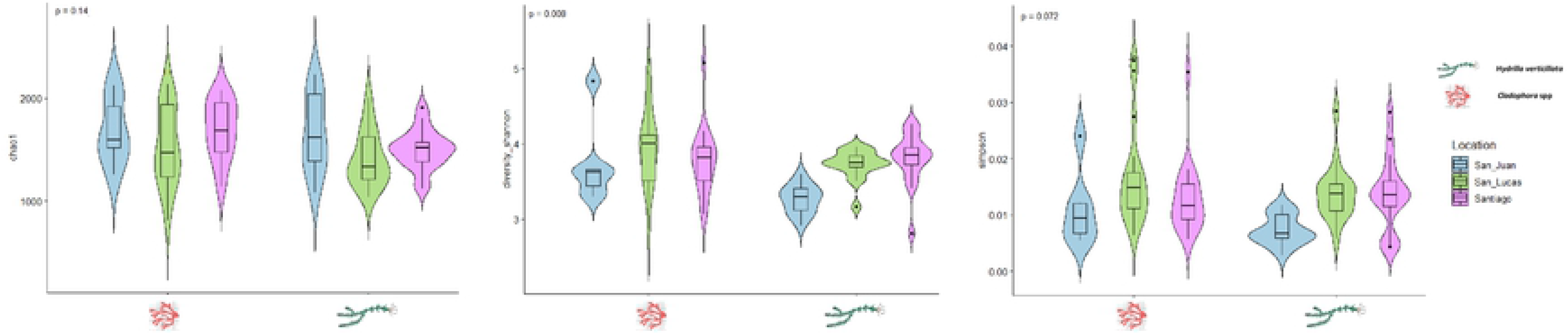
Richness (chao1), diversity indexes (Shannon-Wiener) and evenness (Simpson) in *Hydrilla* and *Cladophora* microbiomes from the three sampling locations.

The result of a slightly different approach, the Linear Discriminant Analysis (Fig.4), highlights microbial taxa most likely to explain differences observed between the microbiomes of *Cladophora* and *Hydrilla* hosts. Fungal genera *Talaromyces* (Ascomycota), *Tylospora* (Basidiomycota), bacterial genera *Trichococcus* (Bacilli), and the cyanobacterium *Chroococcidiopsis* are indicative of the *Cladophora* microbiome. *Hydrilla*-associated microbial indicators are all bacteria: *Exiguobacterium* (Bacilli), *Romboutsia* (Clostridia), *Aeromonas* (Gammaproteobacteria), *Acinetobacter* (Gammaproteobacteria), *Paraclostridium* (Clostridia), and *Clostridium* (Clostridia).

**Figure 4.**
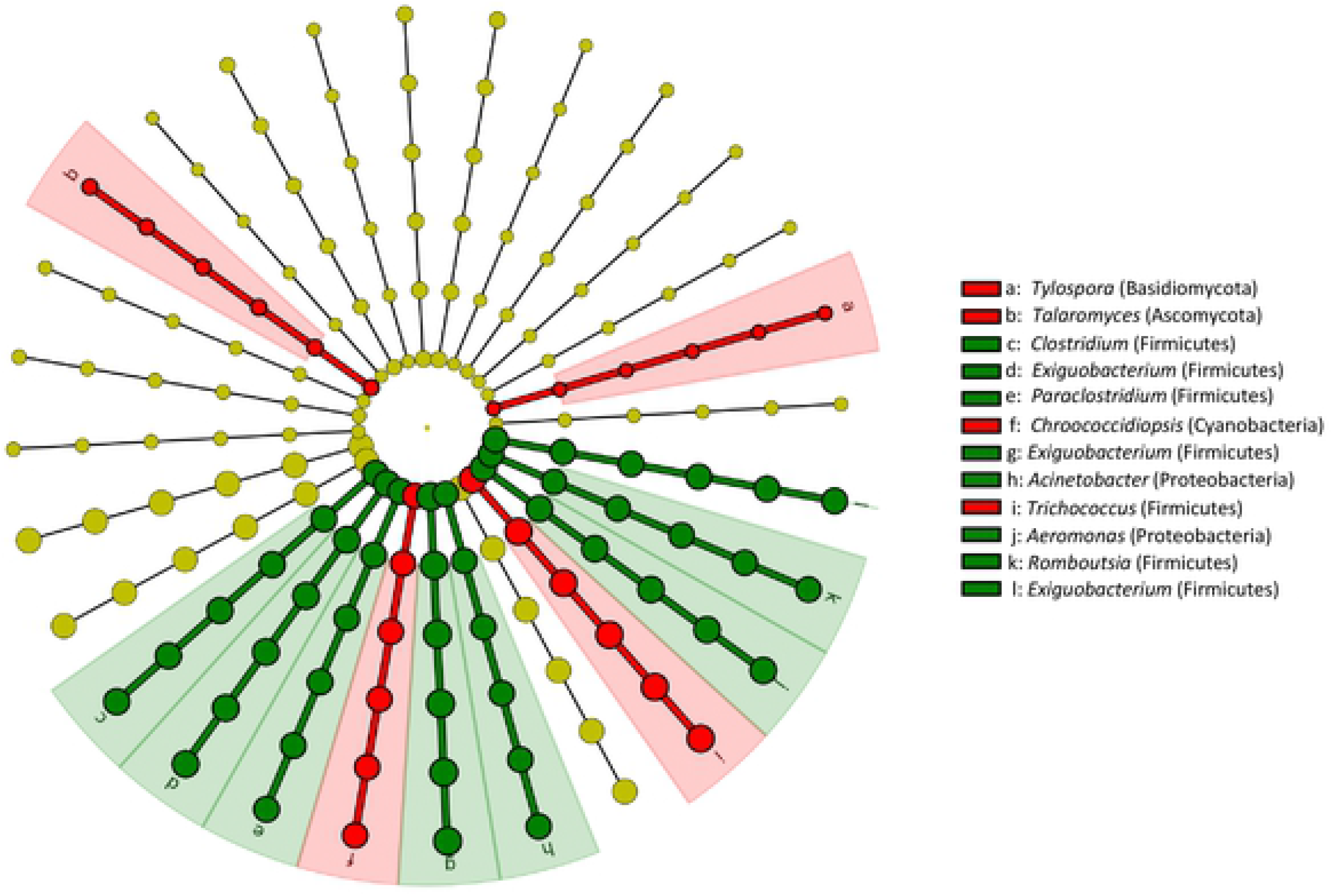
Linear discriminant analysis (LDA) effect size (LEfSe) displaying significant indicator taxa in *Cladophora* (red) and *Hydrilla* (green) microbiomes.

### N_2_-fixation potential and activity

Our dataset contained cyanobacterial sequences, with the average relative abundance 1.4% (± 2.0% SD), ranging between 0.2 and 11% in the different samples (Tab. S1). The *Cladophora* microbiomes contained significantly larger relative proportion of cyanobacterial sequences than *Hydrilla* samples (p = 0.015). The sequence-based taxonomy correlated well with microscopic analysis (Tab.S2). Of the genera capable of N_2_-fixation, both analyses confirmed the presence of the bloom forming filamentous species *Limnoraphis robusta*, and the heterocytous genera *Calothrix* and *Pseudanabaena*. Amplicon sequencing (Tab. S1) revealed the presence of additional N_2_-fixing genera, such as *Chroococcidiopsis*, *Nostoc*, *Scytonema*, *Synechococcus*, and *Tolypothrix*. In addition to diazotrophic cyanobacteria listed above, the amplicon sequencing confirmed the presence of other prokaryotic diazotrophs (Tab. S1). The most important genera, distributed across several bacterial classes, included *Klebsiella*, *Pseudomonas*, *Clostridium*, *Rhodobacter*, *Bacillus*, *Azospirillum*, *Rhizobium*, or *Burkholderia*. Overall, genera known to be able to fix N_2_ consituted roughly 15% on average of all genera found in each sample. *Hydrilla* microbiomes contained slightly but significantly higher number of diazotrophic genera compared to *Cladophora* microbiomes (Fig. 5b), and these genera were also present at a significantly higher relative abundance exceeding 60% of total OTUs on average (Fig. 5a).

**Figure 5.**
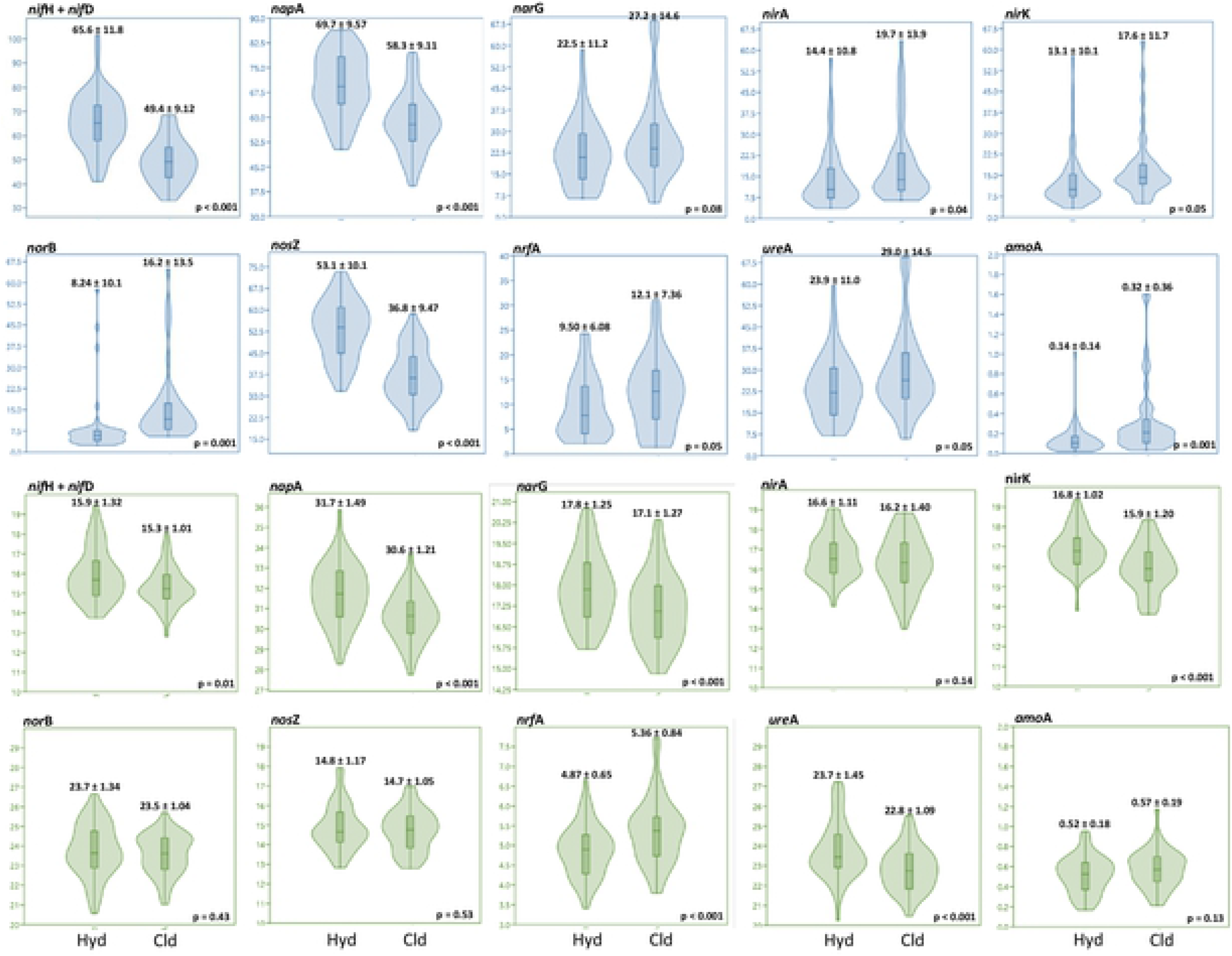
Proportion of functional genes involved in organic matter cycling in *Hydrilla* (Hyd) and *Cladophora* (Cld) microbiomes. The relative abundances of detected taxa are shown in blue (a). Green plots (b) show the percentage (out of the total number of OTUs) of taxa with confirmed presence of the respective functional gene in the genome.

The N_2_-fixation activity, expressed as acetylene-reducing activity of the nitrogenase enzyme (Fig. 6), was highly light-dependent, with daytime activity up to two orders of magnitude higher than nighttime activity in some samples. The difference between daytime and nighttime fixation was significant across all samples, with daytime and nighttime values averaging 45.9 and 8.3 C_2_H_4_ g DW^−1^ h^−1^, respectively. The variability between replicates was high, and no significant difference was found between the nitrogenase activity associated with *Hydrilla* and *Cladophora* across all locations and night and day exposure (mean values of 35.8 and 25.1 nmol C_2_H_4_ g DW^−1^ h^−1^, respectively, Fig. 6). Correlation between the N_2_-fixation activity and the relative proportion of potential N_2_ fixers, both autotrophic cyanobacteria and bacteria, was insignificant for both the *Cladophora* and *Hydrilla*-associated microbiomes (data not shown). The average N_2_-fixation activity values in both *Hydrilla* and *Cladophora* were approximately an order of magnitude lower than those measured in the positive control of *A. filiculoides* (660±26 and 252±76 C_2_H_4_ g DW^−1^ h^−1^ in the “day” and “night” exposures, respectively; n=6). The N_2_-fixation activity associated with *Hydrilla* was comparable to that found in the native macrophytes *P. illinoensis* and *P. pectinatus* from the same location during daytime exposure (Fig. 6). A more pronounced difference was seen during the nighttime exposure, where *Hydrilla*-associated N_2_-fixation activity exceeded that of the other two macrophytes approximately three-fold on average (Fig. 6).

**Figure 6.**
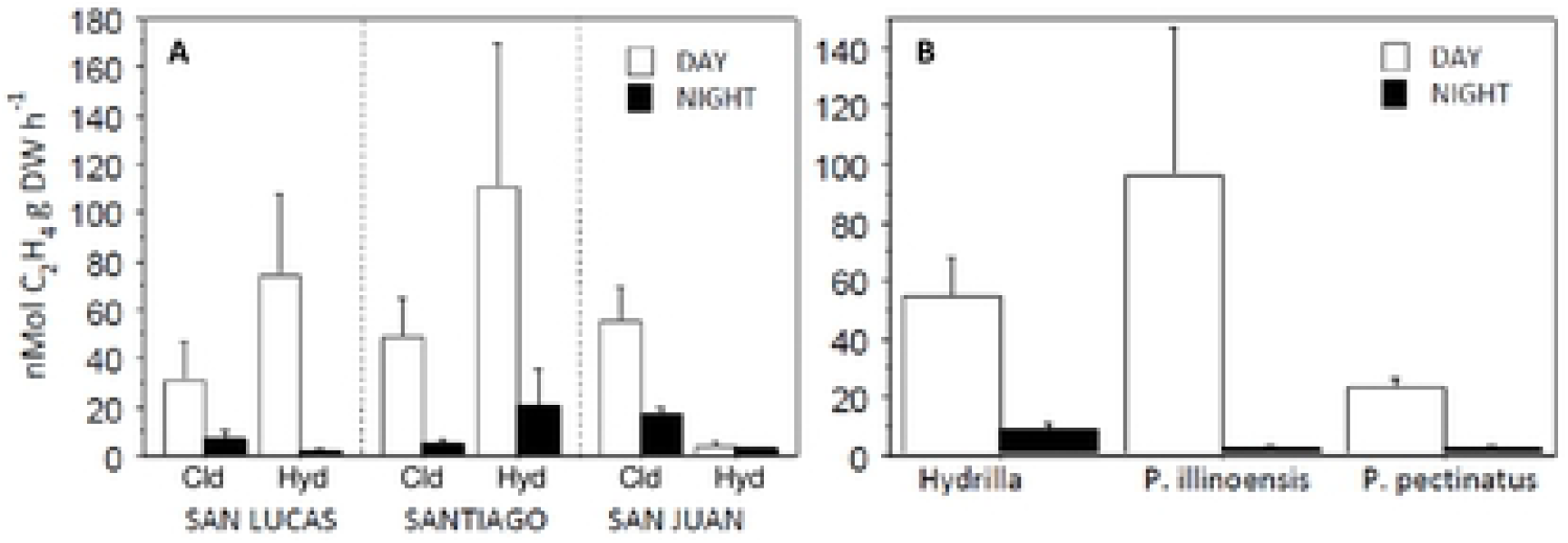
Mean values of N_2_-fixation rate, expressed as C_2_H_4_ g DW^−1^ h^−1^; error bars indicate the standard error of mean, n = 3. A: comparison of locations (San Lucas, Santiago, and San Juan) and microbiomes (Cld = *Cladophora*, Hyd = *Hydrilla*); B: comparison of *Hydrilla* with other two native species *Potamogeton illinoensis* and *P. pectinatus*.

### Nitrogen cycling potential

Regarding other biogeochemical processes related to the N-cycle, denitrification was an activity potentially associated with a high proportion of the total OTUs in most samples (Table S2). On average over a third of the bacterial genera found in the dataset are known to perform the reduction of nitrate to nitrite (*napA*/*narG* marker genes, Fig. 5b), while these constituted a significantly higher relative proportion in the *Hydrilla*-associated microbiomes campared to *Cladophora* samples (Fig. 5a). Capacity to reduce NO_2_^−^ to NO (*nirA*/*nirK* gene markers, Fig. 5b) was previously shown to be present in third of the bacterial genera, while those known to be able to convert nitrous oxide to nitric oxide (*norB* gene marker) were slightly less numerous and less abundant in the datset (Fig. 5b). On average, approximately 15% of bacterial genera present were capable of performing the final step in the denitrification pathway catalyzed by the enzyme N_2_O reductase, and these were significantly more abundant in *Hydrilla* samples (Fig. 5a). Only approximately 5% of bacterial OTUs on average were associated with the direct conversion of nitrite to ammonia (*nrfA* gene marker) and these were significantly more abundant in *Cladophora* microbiomes (Fig. 5a). Genera that are known to convert urea to ammonia (*ureA* marker gene) were also detected, and represented approximately a fifth of the present bacterial taxa on average (Fig. 5b). The relative proportion of ammonia-oxidizing bacteria in the dataset was low, generally below 1% of both total genera counts and relative abundance (*amoA* gene marker, Fig. 5a, b). Some detected fungal taxa have also been implicated in contributing to N-cycling: members of the genus *Rhodotorula* are known to scavenge nitrogenous compounds from its environment remarkably well even at very low concentrations, while the genus *Tylospora* contains known efficient denitrifiers [43].

### Degradation of complex organic matter

Bactrerial genera known to degrade complex organic matter constituted a significant proportion of the dataset. Apart from the typical anaerobic fermenters (such as members of the genera *Clostridium*, *Paraclostridium*, *Romboutsia*, or *Terrisporobacter*), the chitin (*Klebsiella*, *Aeromonas*) and lignin (*Variovorax*, *Burkholderia*) degrading genera (markers chB and ligE, respectively) were detected, the former being significantly more common in the *Cladophora* microbiomes (Fig. 5). Potential to degrade phenolic substances (*laccase* and *ppo* markers) was very common, present in over 25% of the bacterial genera, which were significantly more abundant in *Hydrilla* microbiomes compared to *Cladophora* (Fig. 5). Members of the family *Saprospiraceae,* bacteria able to degrade and utilize complex organic matter, were common in all samples, most of the OTUs only identifiable to the family level (Table S1).

Given the high fungal diversity of the dataset, it is likely that fungi contribute both to complex organic matter mineralisation and production. The dominant fungal groups contain many known degraders of complex compounds, such as Deuteromycetes. Interestingly, The highly abundant yeast-like saprotroph *Aureobasidium pullulans*, typical of the *Hydrilla* microbiomes, produces β-glucan, as well as pullulan (poly-α-1,6-maltotriose), a neutral extracellular polysccharide readily utilised by bacteria [44].

### Growth promotion

The dataset (Table S1) contained a significant proportion of taxa known to positively influence host health or growth by possessing various growth-promoting traits, especially in *Hydrilla*. *Exiguobacterium*, present in the core microbiome, was shown to enhance growth in several plant species by indole acetic acid, siderophore, and hydrogen cyanide production, phosphate solubilization, and antagonism toward pathogens [45]. Similarly, the fungal genera important in all samples *Rhodotorula*, *Aureobasidium*, *Talaromyces*, *Rhizopus*, *Meliniomyces*, *Oidiodendron*, and *Metarhizium* are all plant endophytes with known growth promoting properties [46]. Fungi widely considered as ectomycorrhizal symbionts of terrestrial plants were frequent throughout the dataset, surprisingly commonly associated with the *Cladophora* microbiomes. These include members of the genera *Lactarius*, *Pezoloma*, *Tylospora*, or *Russula* (Table S1).

### Resistance to radiation

There is a conspicuous presence of radiation-resistant bacterial taxa in the dataset, distributed across all samples (Table S1). These include the highly abundant genus *Exiguobacterium*, well as *Deinococcus* spp., *Stenotrophomonas maltophilia*, and *Acinetobacter radioresistens*. One of the most common fungi in the dataset – yeasts belonging to the genus *Rhodotorula* – also synthesize photoprotective compounds [47].

### The presence of potential pathogens

Several bacterial genera that have been implicated as disease causing agents in humans [48], livestock [49], or fish [50] were found in the dataset, some of them at high relative abundances (Table S1). These include *Legionella* (Legionnaires’ disease, Pontiac fever), *Aeromonas* (gastroenteritis and wound infections), *Mycobacterium aubagnense* (pulmonary infections, necrosis), *Enterococcus faecalis* (urinary tract infections), *Escherichia coli* (gastroenteritis, urinary tract infections), *Pseudomonas* (nosocomial infections), *Tsukamurella pulmonis* (bacteraemia, pulmonary infections), *Acinetobacter* (bacteraemia, pulmonary infections, meningitis), *Stenotrophomonas* (emerging multidrug-resistant global opportunistic pathogen), *Flavobacterium* (fish pathogen), *Streptococcus parauberis* (livestock and fish pathogen), or *Kocuria* (*fish pathogen*).

### Co-occurrence analysis

The number as well as the taxonomic profile of the highly connected taxa (i.e. taxa that are considered to play a significant role within the microbiome) were notably different between the *Cladophora* and *Hydrilla* hosts. Co-occurrence network analysis of bacterial and fungal communities consisted of 87 and 36 nodes (bacterial or fungal taxa), 224 and 96 edges (associations among them), and the coefficient of clustering was 0.32 and 0.33 for *Cladophora* and *Hydrilla*, respectively (Fig. 7). The connectedness and the density of the network was higher for *Cladophora* microbiomes, but although it showed higher number of nodes and edges, it also displayed higher heterogeneity, with several disconnected smaller networks, and very few connective nodes between the bacterial and the fungal taxa. The *Hydrilla* microbiome network analysis, on the other hand, resulted in one main network and larger degree of interconnectedness between bacterial and fungal OTUs (for the summary of network parameters see Table S3).

**Figure 7.**
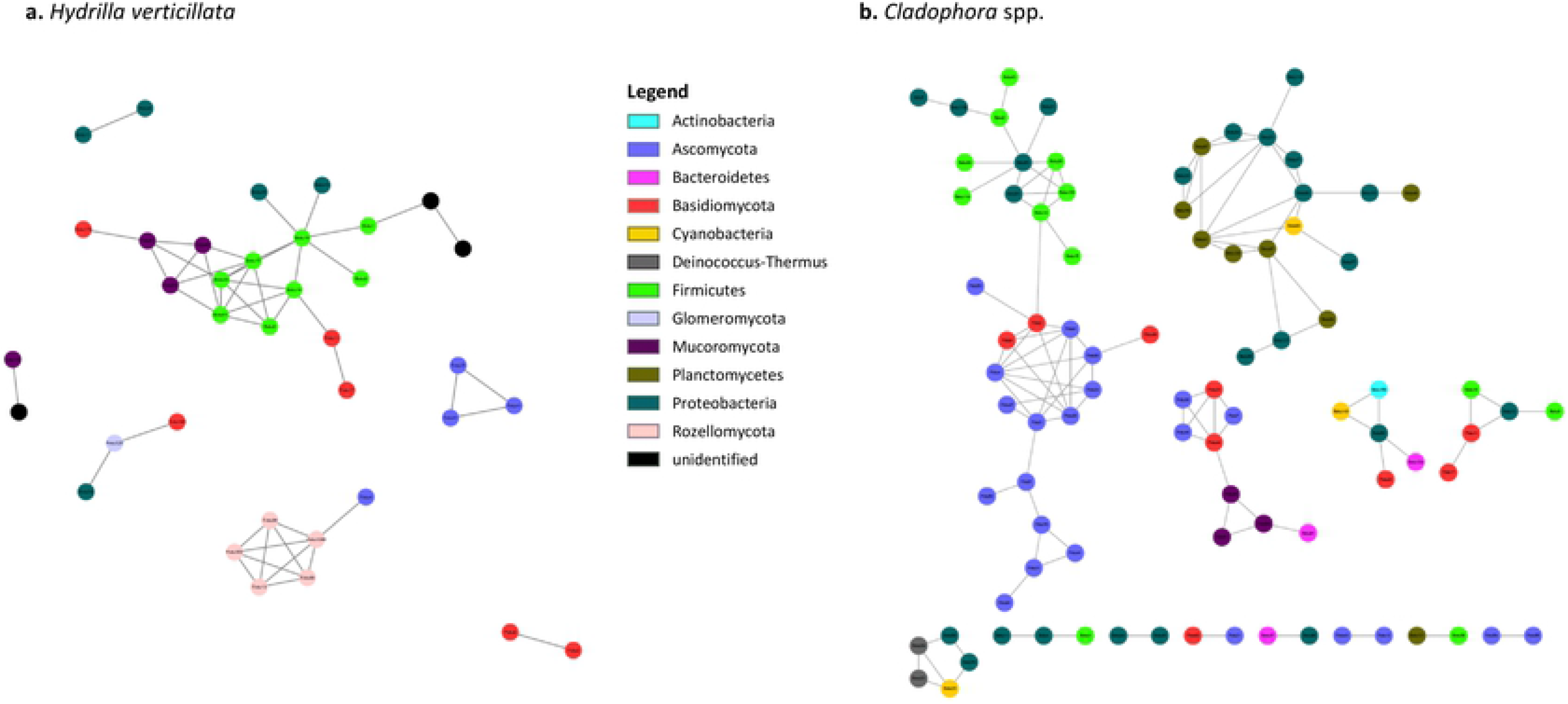
Significant co-occurrence relationships in the microbiomes of a) *Hydrilla verticillata* and b) *Cladophora* spp. hosts. Each node represents a bacterial or fungal OTU, coloured according to taxonomy.

## Discussion

Our dataset clearly shows that both the *Hydrilla* phyllosphere microbiomes and the *Cladophora*-associated microbiomes are highly diverse in terms of bacteria and fungi, and the analysis of putative ecological function using the FunGene pipeline (see Material and Methods) indicates that there is also significant potential for diverse biogeochemical processes and complex intra-microbiome interactions, while the sampling location did not play a significant role in structuring of the microbiomes. Despite some important differences, the very similar structuring and function of microbiomes in *Cladophora* and *Hydrilla* suggest that the same factors govern the interactions between hosts that differ considerably in their phylogeny, anatomy, and physiology and the associated microorganisms.

The roots of submersed plants are greatly reduced in biomass and function compared to the roots of terrestrial plants and their role in nutrient uptake from lake substrates is not well-understood. There is still an ongoing debate in scientific literature whether nutrients in rooted aquatic plants are taken up predominantly from the substrate, the water column, or both. The evidence published so far shows that the mechanisms responsible for nutrient uptake vary greatly among macrophyte species and under different environmental conditions [51]. Published results indicate that the Atitlán native *Potamogeton* species rely more on root nutrient uptake [52] compared to the foliar uptake dominance seen in *Hydrilla* [53]. Since the phyllosphere (here the term also includes leaf-bearing stem parts) likely is the main point of entry of essential nutrients in *Hydrilla*, as well as the main source of photosynthate deposition, its phyllosphere-associated microbiomes likely perform a functional role similar to that of rhizospheric microbiomes of terrestrial plants, including nutrient mobilization.

Looking at the phylum or even family level, the *Hydrilla* and *Cladophora* microbiomes had very similar composition, sharing many core taxa even at the OTU level, despite of the evolutionary distance and anatomical differences between the two host organisms. Many taxa with widely differing life strategy were present in the datasets: bacteria known as strict anaerobic fermenters (for example the Clostridia), the metabolically versatile aerobic *Pseudomonas* spp., the *Phreatobacter* spp. known to be able to grow under conditions of extreme oligotrophy, or members of the typically copiotrophic genera *Bacillus* and *Enterobacter*. Plant endophytes, such as members of the fungal genus *Trichoderma*, and *Oidiodendron*, or the bacterial *Burkholderia* spp. were present in the dataset alongside microbes that likely occupy light-exposed algal filaments and plant leaf surfaces, due to their radiation resistance and photo-protective pigment production traits (see results). Interestingly, an *Oidiodendron maius* strain was shown to upregulate the expression of plant N uptake transporters and thus positively influence N uptake and growth of its host [54].

*Aerobasidium pullulans*, another known fungal endophyte, associated with *Hydrilla*, produces an extracellular polysaccharide pullulan (α-1,4-; α-1,6-glucan) which is known to play a key role in the formation of oxygen impermeable biofilm matrices [55]. This makes *Aureobasidium* one of the prime candidates for an important ecosystem engineering role and together with the other diverse compositional characteristics of the studied microbiomes implies very high niche diversification and the possibility for a wide variety of biogeochemical processes to take place simultaneously.

The presence of potential pathogens in the dataset likely reflects the known ability of many macrophytes to significantly reduce pathogen loads in wastewaters [56].

The capability of *Hydrilla* to decrease the amount of coliforms in waste water by up to 95% was documented by [57] and our unpublished data from Atitlán show similar efficiency for both the total coliforms and *Escherichia coli*, 95% and 100%, respectively. *Hydrilla* thus likely serves as an effective water sanitizing filter, however, more data are needed to further test this hypothesis and clarify the mechanisms involved.

With regards to our research question of whether the associated microbes can contribute to N acquisition in Hydrilla, we found a relatively high proportion of N_2_-fixing genera, both cyanobacteria and other bacteria, in the dataset and confirmed their activity experimentally, showing that the process also takes place during nighttime. This is what distinguished *Hydrilla* from the native *Potamogeton* spp., and, as N_2_-fixation is a carbon-limited process, may be related to the fact that *Hydrilla* is able to fix carbon into malate and aspartate during a nighttime C4-like metabolism. The high potential for organic matter mineralization apparent in the dataset may be another indication of possible mineral N source for the plants. The genus *Pirellula* (Planktomycetacia) as well as other members of the class were some of the most important taxa, ubiquitous in all samples at high proportions of total reads (TableS1). Common in phytodetrital macroaggregates, they are known to catalyze the initial aerobic breakdown of complex organic polymers into simpler compounds, thereby performing a key step in the C and N cycles [58]. The potential ability to degrade complex polymers however, was associated with large proportions of total OTUs across all our samples. One of the most abundant and widespread polymers in aquatic environments is chitin, as it serves as a structural element in many organisms (e.g. fungi, crustaceans, or algae) [59]. The cell walls of *Cladophora* also contain a significant outer layer of chitin. The high abundance of chitinolytic genera in the dataset in general, but specifically the significantly higher proportion found associated with *Cladophora* compared to *Hydrilla* (Fig. XY), suggest that the decomposition of chitin is an important process in the microbiome of these hosts, and, because of the high N content of chitin (up to 7%), may also contribute significantly to N-cycling within the algae-plant-microbe system [60]. While it has also been shown that many soil fungi can use chitin as a sole source of N (e.g. Leake & Read 1990), whether aquatic fungi can also tap into this abundant N-source needs further study.

Apart from the chitinolytic taxa, many additional bacterial genera that were previously shown to have aromatic compound degrading ability and to use polyphenol oxidases, laccases, or beta-etherases to access nutrients in polymers such as lignocellulose, lignin, or phenolic compound-containing humic substances, were present in the dataset in high relative abundances, especially in *Hydrilla*-associated microbiomes, as these substances are primarily associated with plant biomass (see Fig. 5 and Table S2). Because of this large potential, further confirmed by the high biomass turn-over rates in *Hydrilla* stands measured by Rejmánková et al. 2018b, it is possible that *Hydrilla* and *Cladophora* microbiomes are involved in some form of „N mining”. It has been shown previously on decomposing leaf litter in soil, that low N availability can actually increase litter decomposition as microbes use labile substrates to acquire N from recalcitrant organic matter. Considering the continuously rising trophy levels in lake Atitlán and the fact that the „N mining” process is consistently suppressed by high N supply or substrate N concentrations [62], assessing the accompanying changes in the functional composition of the microbiomes and the associated alterations in the algae-plat-microbe interactions merits further study.

The presence of several typically ectomycorrhizal fungal genera associated with both *Cladophora* filaments and *Hydrilla* leaves was surprising to us and we considered the possibility of sample contamination by spores being wind-blown onto the lake water surface from the shore. This is however unlikely given the consistency of the pattern. Aquatic fungi are an overlooked and highly understudied group, despite its potential importance in aquatic food-web ecology [63]. Furthermore, there are studies which confirm the presence of ectomycorrhizal species, such as *Russula*, in aquatic environments [64]. It remains to be determined whether the association between these fungi, filamentous algae, and macrophyte leaves represents a growth advantage.

In addition to the processes that could supply N into or recycle it within the system, our data indicate that denitrification is an ability potentially present in a large number of bacterial and fungal taxa in the dataset, and, if taking place, may represent important means of N loss from the littoral ecosystem. While the results of some studies of the macrophyte beds show positive net N fluxes due to efficient N_2_-fixation [65], other studies on peri/epiphytic communities report net N loss by denitrification [66]. The outcome of the N_2_-fixation (source)/denitrification (sink) coupling is clearly dependent on many factors, including nitrate availability in the system, dissolved organic carbon availability, season, or dissolved oxygen. As we have only measured N_2_-fixation activity, we cannot say anything about the net flux of N in the Atitlán littoral communities, although significant rates of epiphytic denitrification are frequently found associated with submersed vegetation despite the high oxygen concentrations in the surrounding water [67]. However, the importance of perifytic denitrification is only likely to increase with increasing lake trophy and, therefore, given the current eutrophication rates in the lake watershed, the stands of the invasive *Hydrilla* hosting denitrifying microorganisms may play an increasingly important role in removing excess N from the watercolumn of the littoral zones in the near future.

### Theoretical N budget in the Atitlán littoral areas with the presence of Hydrilla stands

Based on the results of this study, as well as previously published data, we attemted to estimate the contribution of N_2_-fixation associated with *Hydrilla* phyllosphere microbiomes to covering the N metabolic needs of the host plant. Based on lake Atitlán bathymetry [26], the area of the littoral zones capable of supporting *Hydrilla* growth is 4,86 km^2^. We estimate that approximately half of that area is covered by *Hydrilla* stands. According to published results [2], the average biomass of *Hydrilla* leaves that are exposed to photosynthetically active radiation is 600 g m^−2^, which represents 1. 458. 000 kg of leaf DW in the littoral zone. According to our average measured N_2_-fixation rates (assuming that 3 mol of C_2_H_4_ produced represent 1 mol of N_2_ reduced, and that there are 200 days per year suitable for N_2_-fixation), the process can supply 2.2 g N kg^−1^ DW day^-1^ into the system, which represents 3.2 tons of N littoral^−1^ year^−1^. *Hydrilla* biomass contains approximately 2% of N [2]. Because of the missing seasonality, the *Hydrilla* whole-shoot biomass (leaves and stems) in the lake littoral can be estimated at 3000 tones throughout the year. If we assume a biomass turnover time of 4 months [2] and 40% N reutilisation efficiency (as estimated for submersed, fast growing macrophytes by Adamec 2014), then the *Hydrilla* stands contain approximately 110 tons of N, of which at least 3% can be supplied by microbial N_2_-fixation alone (likely a conservative estimate). However, the presence of efficient microbial decomposers, the very fast rates under which the senescent biomas is turned over [2], and the fact that *Hydrilla* stands represent barriers which effectively retard water flow and exchange [11,69] are likely responsible for the elevated nutrient levels observed in littoral watercolumn compared to the pelagial (Table 1). While we have very limited information on the mechanisms of N loss from the system, such as microbial denitrification, and no quantitative data, it can be assumed that a significant proportion of the N liberated by decomposition is availble for uptake by the growing *Hydrilla* shoots.

## Conclusions

Numerous studies have considered invasive aquatic plants with regards to their growth, regeneration capacity, photosynthesis traits, genetic, reproductive, overwintering strategies and management, however, very little attention has been given to the link between the associated microbial community structure and function, and invasive plant success in a given ecosystem [25]. In this respect the research of macrophyte invasions has lagged behind that in terrestrial plants, where studies, based mostly on the rhizosphere-associated microbes, clearly show that microbiomes can be instrumental in facilitating plant invasions [70]. We conclude that phyllosphere microbiomes of fast growing submerged macrophyte have the compositional and functional diversity that allows them to influence not only the successful establishment of invasive species, but also the biogeochemical cycling of entire shallow lake ecosystems.

## Acknowledgements

The study was supported by the Grant Agency of the Czech Republic project no. P504/17-10493S. We thank the Asociación de Amigos del Lago de Atitlan, CEA–UVG Altiplano, and AMSCLAE for logistical and personnel support. We would also like to thank Martina Čtvrtlíková and Lubomír Adamec for their valuable comments during the manuscript writing period.

## Supporting material

**Figure S2.**
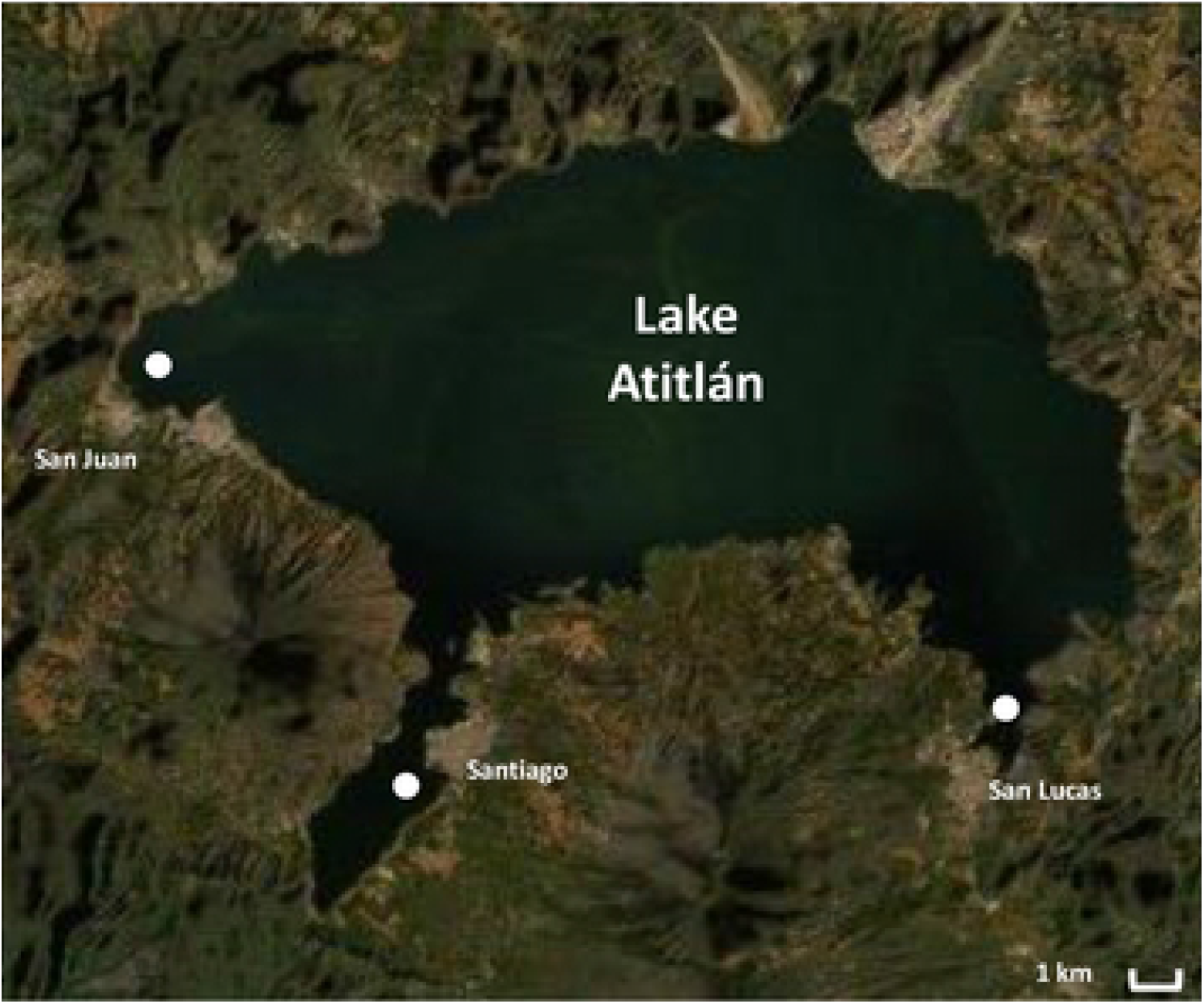
Lake Atitlán, Guatemala, CA, with sampling locations (filled circles) near the San Juan, Santiago, and San Lucas settlements (Image source: Scribblemaps.com, 2020).

**Table S1.** Operational Taxonomic Unit (OTU) table listing all bacterial and fungal OTUs identified in the study.

**Table S2.** Functional predictions for bacterial taxa (FunGene database output)

**Table S3.** Co-occurrence analysis results listing important network parameters

**Table S4.** Mapping file containing relevant metadata

